# Notch Signaling Commits Mesoderm to the Cardiac Lineage

**DOI:** 10.1101/2020.02.20.958348

**Authors:** Evan S. Bardot, Bharati Jadhav, Nadeera Wickramasinghe, Amélie Rezza, Michael Rendl, Andrew J. Sharp, Nicole C. Dubois

## Abstract

During development multiple progenitor populations contribute to the formation of the four-chambered heart and its diverse lineages. However, the underlying mechanisms that result in the specification of these progenitor populations are not yet fully understood. We have previously identified a population of cells that gives rise selectively to the heart ventricles but not the atria. Here, we have used this knowledge to transcriptionally profile subsets of cardiac mesoderm from the mouse embryo and have identified an enrichment for Notch signaling components in ventricular progenitors. Using directed differentiation of human pluripotent stem cells, we next investigated the role of Notch in cardiac mesoderm specification in a temporally controlled manner. We show that transient Notch induction in mesoderm increases cardiomyocyte differentiation efficiency, while maintaining cardiomyocytes in an immature state. Finally, our data suggest that Notch interacts with WNT to enhance commitment to the cardiac lineage. Overall, our findings support the notion that key signaling events during early heart development are critical for proper lineage specification and provide evidence for early roles of Notch and WNT during mouse and human heart development.

**Summary statement:** Early fate decisions are dictated by the embryonic signaling environment. We show that Notch signaling is active during early mouse development and that activating Notch in human cardiac mesoderm enhances cardiomyocyte differentiation efficiency.

## Introduction

An anatomically and functionally normal heart is the result of a coordinated series of specification events and cell movements that begin as early as gastrulation (Meilhac et al., 2014; Vincent and Buckingham, 2010). Fate mapping of the early mouse embryo revealed the organized manner in which cells of the epiblast migrate through the primitive streak (PS) to become members of the separate germ layers (Lawson et al., 1991). The consistency of these movements strongly suggests that the location and timing of ingression through the PS are determinants of cell fate. Notably, the earliest cells to ingress migrate across the embryo to become anterior structures, with those that do so later taking up more posterior positions (Lawson et al., 1991). This pattern is consistent in zebrafish, with the first cells to gastrulate migrating to the future heart region ahead of those that ingress later (Keegan, 2004). A series of experiments in avian embryos support the findings from mice, which assigned the prospective cardiogenic region of the presumptive PS and further delineate the time sensitivity of gastrulation movements in chick heart development (Garcia-Martinez and Schoenwolf, 1993; Inagaki et al., 1993). Importantly, epiblast cells fated to contribute to the heart are not committed to the cardiac lineage until shortly after they ingress through the PS, suggesting that lineage commitment occurs after gastrulation (Inagaki et al., 1993; Tam et al., 1997; Yutzey and Bader, 1995).

Recent studies using genetic lineage tracing approaches have greatly advanced our understanding of these earliest events during heart development. Studies using *Mesp1-Cre* lineage tracing have further solidified the mechanisms proposed from fate mapping data (Devine et al., 2014; Lescroart et al., 2014; Saga et al., 1999). Specifically, the first cells to express *Mesp1* contribute to derivatives of the first heart field, primarily the left ventricle. As more cells express *Mesp1* and become specified to the cardiac lineage, they contribute to the heart from the right ventricle, atria, and finally outflow tract (Lescroart et al., 2014). Previous work from our lab revealed the presence of a population of cells that transiently expresses *Foxa2* during gastrulation and gives rise specifically to ventricular cardiovascular cells of the heart, likely representing cells that ingress and express *Mesp1* early (Bardot et al., 2017b). The striking similarity between the mouse, zebrafish, and chick models suggest conservation of the regional organization and sequence of early events during heart morphogenesis. The tools developed in these studies enable us to understand the early steps of heart development with increased resolution, at the time when heart progenitors are becoming committed to the cardiac lineage.

While patterns of early migration are well established, the heterogeneity within the cardiac progenitor pool and the signaling environments to which early and late-specified cells are exposed are not yet well defined. Recent work has dissected the transcriptional landscape within the *Mesp1*-expressing cardiac progenitor pool (Lescroart et al., 2018; Scialdone et al., 2016). These analyses showed that cardiac progenitors are largely homogenous through E7.25. Between E7.25 and E7.5, as cells progress along the differentiation trajectory, two groups can be identified reflecting the first and second heart field lineages (Lescroart et al., 2018). What remains unclear is how exposure to different signaling environments affects the differentiation trajectories of these progenitor populations.

Here, we have isolated several distinct mesoderm and endoderm populations from the mid gastrulation mouse embryo and utilized comparative transcriptome analysis to identify key regulatory pathways characteristic of each subpopulation. Specifically, we identified Notch signaling as a pathway for *in vivo* cardiac mesoderm specification, and showed that transient induction of Notch signaling in human pluripotent stem cell (hPSC)-derived mesoderm enhances cardiomyocyte differentiation. Cardiomyocytes from Notch-induced cultures were less mature than their control counterparts and lacked a clear atrial vs. ventricular identity. We show that the increase in cardiomyocyte generation is a mesoderm specific effect, likely through interaction with other pathways important for mesoderm specification. We further validated this hypothesis through the modulation of

Notch and WNT signaling in hPSC differentiation to cardiomyocytes, demonstrating that Notch signaling activation is sufficient for commitment to the cardiac lineage and necessary for differentiation in the absence of WNT inhibition. These data help us to better understand the earliest steps during cardiac commitment and highlight the similarities between the mouse and human differentiation programs.

## Results

### Identification and characterization of distinct mesoderm subpopulations in the gastrulating mouse embryo

We have previously used lineage-tracing with *Foxa2Cre* to identify a cardiac progenitor population that gives rise selectively to the ventricles of the heart, but not the atria (Sup. Fig. 1a) (Bardot et al., 2017b). To label the ventricular lineage during embryonic development, we crossed *Foxa2Cre* mice with *ROSA26-mTmG* reporter mice, which express membrane-tethered Tomato in cells unexposed to Cre, but express membrane-tethered GFP after Cre recombination. *Foxa2* lineage-traced cells (mGFP+) can be identified within the migratory mesodermal wings shortly after ingressing through the primitive streak (Sup. Fig. 1b), and later in the cardiogenic region below the head folds at the anterior side of the embryo (Sup. Fig. 1c). *Foxa2* lineage-traced cells are restricted to a subdomain of the cardiac crescent and localize to the ventricular regions of the heart at all stages examined (Sup. Fig. 1d-g). This approach therefore provides us the unique opportunity to examine the ventricular and atrial lineages at any stage during development, starting as early as during gastrulation.

To characterize the earliest stage of such lineage specification we focused on mid gastrulation (E7.25), shortly after cardiac mesoderm is formed. At this point during development, ventricular cardiac mesoderm (vCMes) migrates anteriorly, just ahead of the atrial cardiac mesoderm (aCMes) (Fig. 1a). Whole mount immunofluorescence (WMIF) analysis of Foxa2Cre:ROSA26-YFP embryos revealed Foxa2Cre:YFP+ cells that express the mesodermal marker PDGFRA as well as Foxa2Cre:YFP-PDGFRA+ cells (Fig. 1b). To elucidate the mechanisms at play as the different mesoderm populations migrate to the anterior side of the embryo and become progressively specified, we FACS purified cells from pooled, stage-matched embryos of the same litter for whole transcriptome RNA sequencing (Fig. 1c). We used antibodies population (Sup. Fig. 2b). against the cell surface markers PDGFRA and KDR to isolate CMes from the Foxa2Cre:YFP+ and Foxa2Cre:YFP-lineages, representing early migrating vCMes and late migrating aCMes, respectively (Fig. 1d, e) (Bardot et al., 2017b; Katt-man et al., 2011). We also isolated hematopoietic mesoderm (HMes, KDRhigh) and endoderm cells (End, Foxa2Cre:YFP+, EPCAM+) as a non-meso-dermal reference (Fig. 1f, g). From pooled litters, we obtained on average 273 vCMes, 840 aCMes, 936 HMes, and 1536 End cells (Sup. Fig. 2a). The difference in the number of each CMes subset recovered reflects their proportion of the total CMes

**Figure 1:**
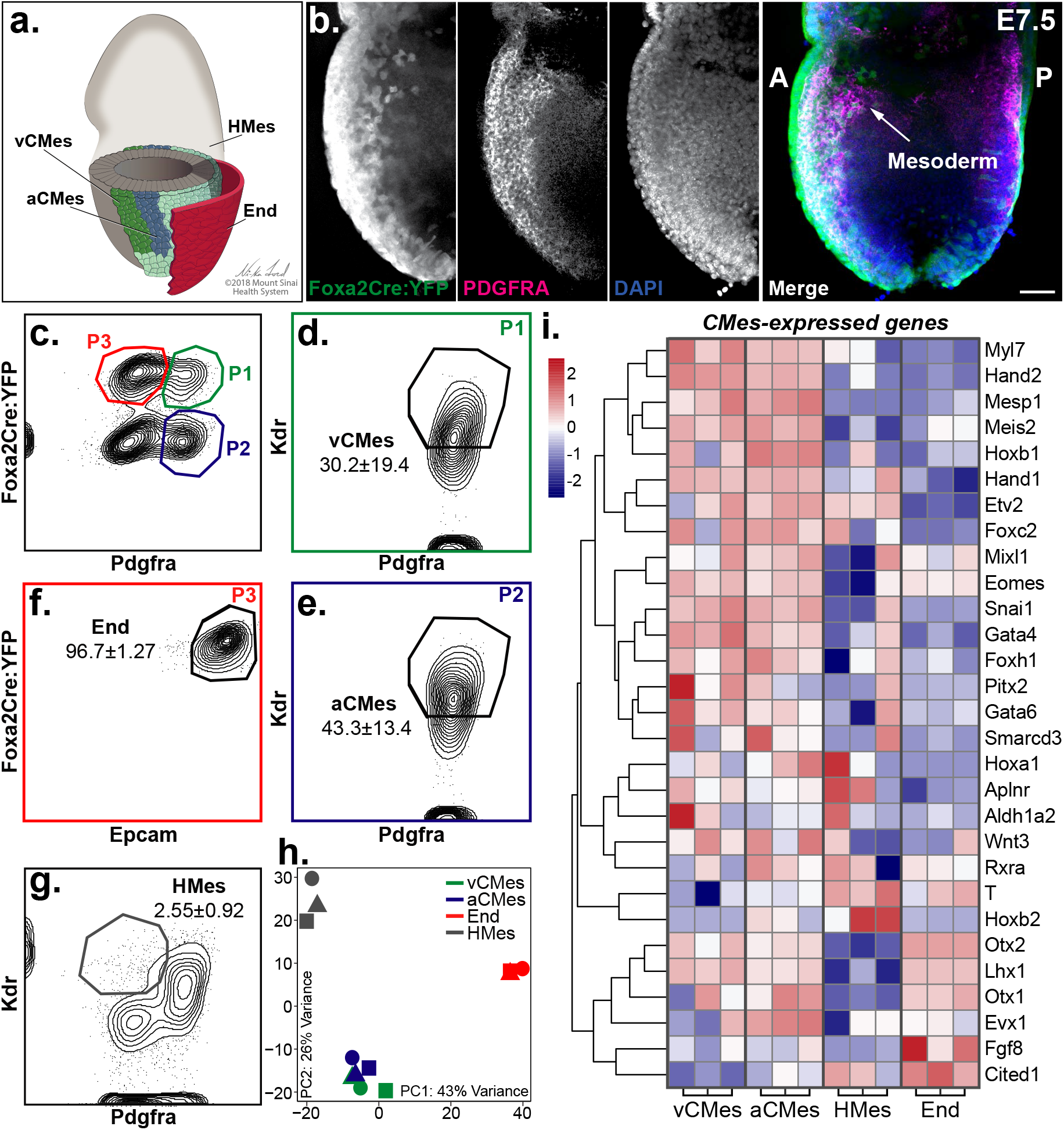
Isolation and gene expression profiling of early progenitor populations from mouse embryos. (a) Schematic of E7.5 embryo and populations of interest, including ventricular and non-ventricular cardiac mesoderm, endoderm, and hematopoietic mesoderm. (b) WMIF analysis of an E7.5 *Foxa2Cre:YFP* embryo with antibodies against YFP and PDGFRA. (c) Isolation strategy for fluorescence activated cell sorting from pooled *Foxa2Cre:YFP* embryos. Cells are first separated by expression of PDGFRA and YFP. (d) Ventricular (YFP+, P1) and (e) non-ventricular (YFP-, P2) CMes are identified by expression of PDGFRA and KDR. (f) Endoderm (YFP+, P3) is identified by expression of EPCAM. (g) Hematopoietic mesoderm (Hem. Mes.) is isolated based on single positivity for KDR. (h) Principal component analysis of whole-transcriptome RNA-sequencing data from isolated populations. (i) Expression z-scores of selected genes known to be expressed in Mesp1+ cardiac progenitors (genes taken from (Lescroart et al., 2014)). A, anterior; P, posterior. Scale bar is 100μm.

Three biological replicates passed a number of QC steps, including hierarchical clustering after batch correction for hidden covariates (Sup. Fig. 2c, d). Faithful expression of signature genes for cardiac mesoderm, endoderm, and hematopoietic mesoderm confirmed the expected overall identity of the isolated cell populations (Sup. Fig 2e). Principal component analysis (PCA) separated the mesoderm and endoderm lineages (PC1, 43% variance captured), and the cardiac from non-cardiac samples (PC2, 26% variance captured) (Fig. 1h). As expected, PCA shows a high similarity between the different CMes populations at this stage, as these share the core cardiac mesoderm gene signature (Fig. 1h). Both vCMes and aCMes samples expressed genes known to be enriched in Mesp1+ cardiac progenitors (Fig. 1i) (Lescroart et al., 2014).

We have previously performed gene expression analysis on mouse embryonic stem cell-derived CMes populations representing the Foxa2+ and Foxa2-progeny *in vitro* (Bardot et al., 2017b). PCA comparing the *in vitro* populations with their *in vivo* counterparts initially showed large differences between the two systems, with PC1 separating the two datasets (Sup. Fig. 3a). Analysis of additional components restored the biological lineage relationships observed previously. Specifically, PC2 separated the mesoderm and endoderm populations, and PC3 isolated the cardiac mesoderm from non-cardiac lineages (Sup. Fig. 3a, b). Direct comparison of PC2 and PC3 resembled the relationships seen when analyzing either *in vitro* or *in vivo* samples individually (Sup. Fig. 3c). Examining the genes driving the variance for each PC reveals that characteristic marker genes for mesoderm, endoderm, or cardiac mesoderm account for the separation seen in PC2 and PC3 (Sup. Fig. 3d).

To elucidate differences between the vCMes and aCMes populations we first performed differential gene expression analysis. We found 416 differentially expressed genes (FDR<5%, fold change>1.5), with 129 and 287 genes enriched in the vCMes and aCMes populations, respectively (Fig. 2a). We focused our analysis on signaling pathways since we hypothesize that the signaling environment at this stage of development will affect cell fate decisions. Of the top 200 most up-and down-regulated genes, 26 have been shown to have signaling roles (Fig. 2b). Further analysis using KEGG pathway datasets revealed Notch signaling to be the only statistically enriched pathway within the vCMes population (Fig. 2c). This is consistent with our previous *in vitro* CMes gene expression data, illustrating the comparability of the *in vivo* and *in vitro* systems at similar stages of differentiation (Fig. 2c) (Bardot et al., 2017b). In addition, active Notch signaling could be observed in E7.5 mouse embryos, as detected by whole mount immunofluorescence for the Notch Intracellular Domain (NICD) (Fig. 2d). NICD+ nuclei are found within the PDGFRA+ mesoderm and can be seen in both the ventricular (Foxa2Cre:H2B-mCherry+) and atrial (Foxa2Cre:H2B-mCherry-) lineages. Overall, these analyses show that Notch signaling is active within the cardiac mesoderm *in vivo* and suggest a temporal gradient in the induction of this pathway during CMes differentiation.

**Figure 2:**
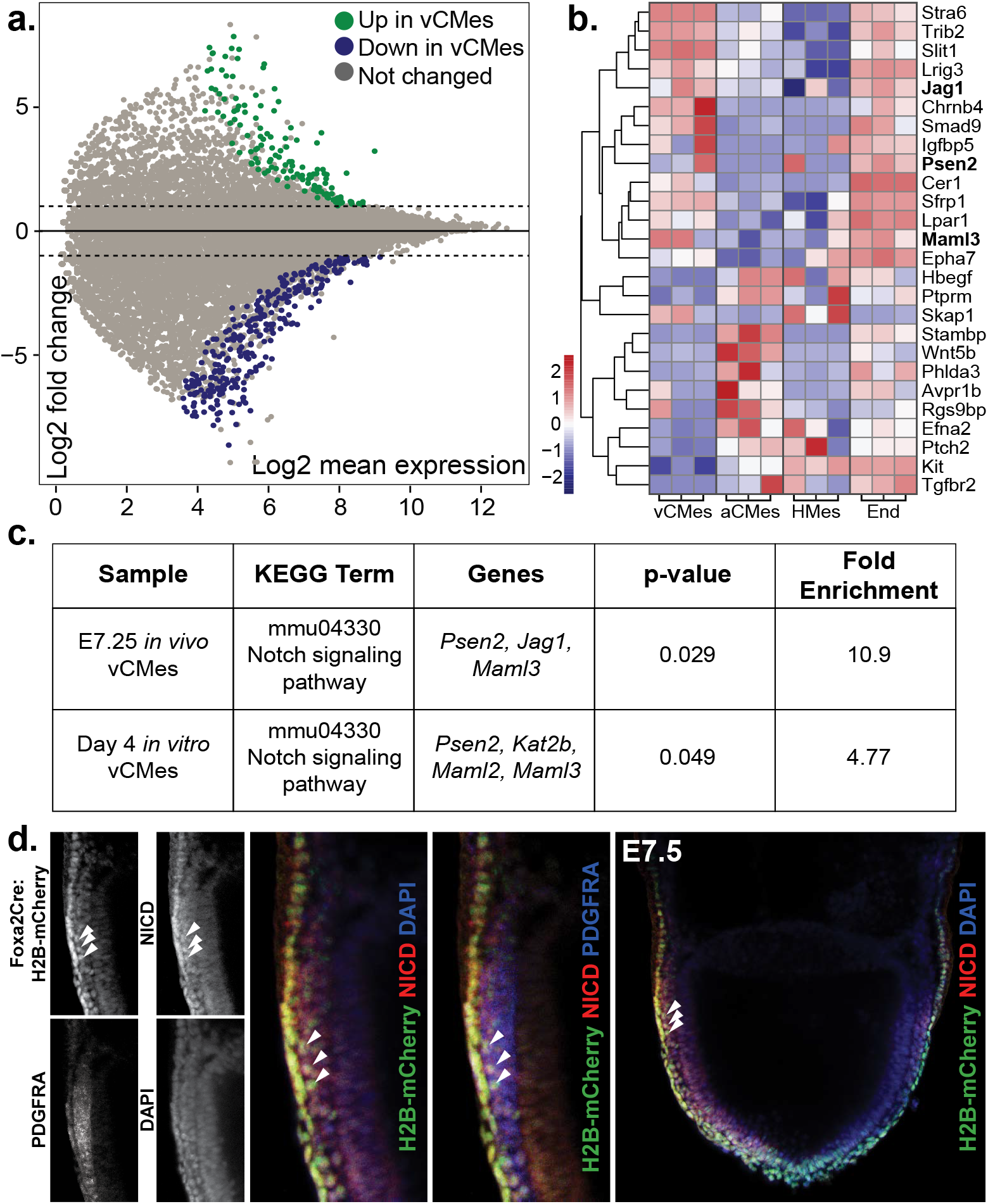
Gene expression analysis reveals a potential role for Notch signaling during cardiac mesoderm differentiation. (a) MA plot showing genes differentially expressed (FDR 5%, fold change >1.5) between *in vivo* YFP+ CMes and YFP-CMes. (b) Expression z-scores for differentially expressed signaling pathway components. (c) KEGG pathway analysis for genes enriched in YFP+ CMes *in vivo*, and comparison with *in vitro* cardiac mesoderm data. (d) WMIF analysis of an E7.5 *Foxa2Cre:H2B-mCherry* embryo with antibodies against mCherry, NICD, and PDGFRA. Embryo is imaged en face to show bilateral mesoderm.

### Activation of Notch signaling enhances cardiomyocyte differentiation from human pluripotent stem cells

We have identified Notch pathway genes as being differentially expressed in cardiac mesoderm populations in mice, suggesting a potential role during early differentiation. To dissect the role of Notch during cardiac mesoderm commitment and differentiation in a temporally-controlled manner, we turned to the well-established human pluripotent stem cell (hPSC) differentiation system. In this model, hPSCs are differentiated in the presence of signaling molecules that recapitulate the *in vivo* signaling environment (Kattman et al., 2011; Yang et al., 2008). In doing so, cells are guided along a normal developmental trajectory first exiting pluripotency, being patterned into mesoderm and specified to cardiac mesoderm, becoming committed to the cardiac lineage, and finally differentiating into functional cardiomyocytes (Fig. 3a) (Calderon et al., 2016; Kattman et al., 2011; Lian et al., 2013; Yang et al., 2008). To specifically manipulate Notch signaling during this process, we used an hPSC line that expresses a Notch intracellular domain-estrogen receptor (NICD-ER) fusion from the human ROSA26 locus (Ditadi et al., 2015). These cells ubiquitously express the NICD-ER protein, which can be activated through addition of 4-hydroxytamoxifen (4OHT) to allow translocation to the nucleus and activation of Notch target genes.

**Figure 3:**
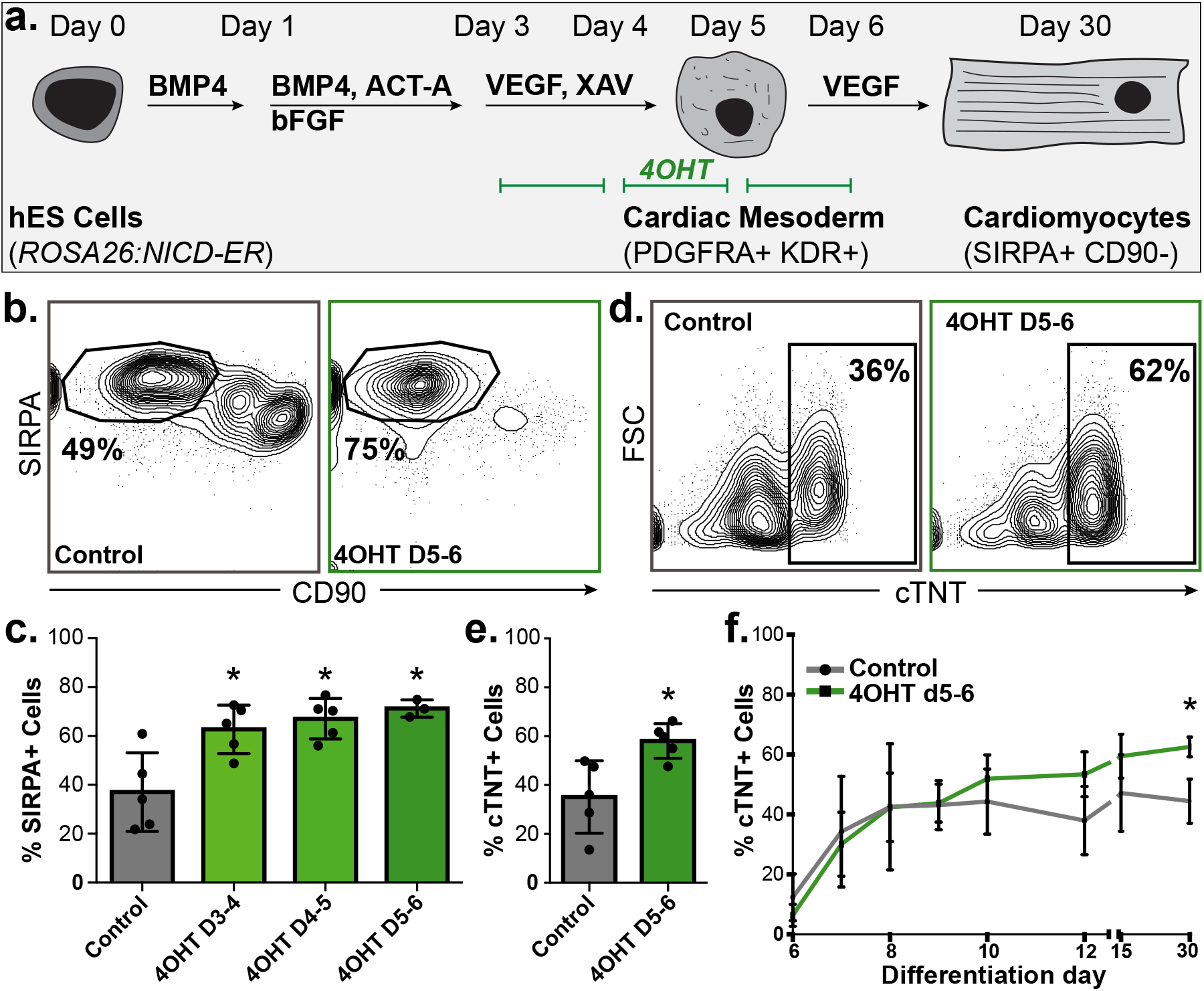
Activation of Notch signaling enhances cardiomyocyte differentiation from human pluripotent stem cells. (a) Schematic of *ROSA26:NICD-ER* hPSC-cardiomyocyte differentiation and Notch induction times. (b) Flow cytometric analysis with antibodies against SIRPA and CD90 to measure cardiomyocyte (SIRPA+CD90-) differentiation efficiency at day 30 in control cultures (left) or after 4OHT treatment from day 5-6 (right) (c) Quantification of data in (b). (d) Flow cytometric analysis of day 30 control (left) or Notch-induced (right) cultures with antibodies against cTNT. (e) Quantification of data in (d). (f) Flow cytometric measurement of cTNT expression over the course of differentiation (n=3 differentiations).

Given our hypothesis that Notch signaling impacts cardiomyocyte differentiation during the specification or commitment of early mesoderm, we first determined the effect of Notch induction during early hPSC cardiomyocyte differentiation. 4OHT was added for 24-hour pulses between day 3 and day 6 of differentiation and cardiomyocyte differentiation efficiency was assessed at day 30. Control cultures yielded 37±16% cardiomyocytes at day 30, as measured by the expression of SIRPA and absence of CD90 (Fig. 3b, left panel) (Dubois et al., 2011). Induction of Notch signaling from days 3-4, 4-5, or 5-6 increased the efficiency of differentiation, with 71±3.6% cardiomyocytes being generated after Notch induction from day 5-6 (Fig. 3b, right panel, quantified in c). Consistent with the data obtained from the *in vivo* CMes RNA-seq experiments, the strongest effect was seen upon Notch induction at day 5 of differentiation when CMes corresponding to E7.25 mesoderm is present *in vitro*. We confirmed the increase in cardiomyocyte generation using flow cytometry for the bona fide cardiomyocyte marker cardiac troponin T (cTNT) (Fig. 3d, quantified in e). Furthermore, we showed that the differentiation kinetics are not significantly affected upon Notch induction, as the onset of cTNT expression occurs at the same time in control and 4OHT-treated cultures (Fig. 3f).

To determine if Notch is required for cardiomyocyte differentiation under these conditions, we also differentiated cells in the presence of the gamma-secretase inhibitor DAPT, which blocks cleavage of the Notch receptor, thereby inhibiting endogenous signaling activity (Sup. Fig. 4a) (Dovey et al., 2001; Morohashi et al., 2006). Treatment with DAPT for the same 24h time periods had no effect on cardiomyocyte differentiation overall (Sup. Fig. 4b, c). To ensure that this observation was not due to the limited time of inhibition or leakiness of the NICD-ER construct, we performed additional inhibition experiments using a human induced PSC (hiPSC) line. We differentiated hiPSCs in the presence of DAPT from days 3-6, covering the key early mesoderm formation and specification events (Sup. Fig. 4d). These experiments confirmed that inhibition of Notch signaling has no effect on cardiomyocyte differentiation efficiency overall (Sup. Fig. 4e, f), and is consistent with recent reports showing no effect from DAPT treatment in standard differentiation conditions (Biermann et al., 2019). This suggests that either Notch signaling is not required for cardiomyocyte differentiation, or that the culture conditions provide a signaling environment that is sufficient to generate cardiomyocytes in the absence of active Notch signaling.

### Cardiomyocytes derived from Notch-induced cardiac mesoderm are developmentally immature

Key signaling events during early differentiation have been shown to impact cardiomyocyte subtype specification, both *in vivo* and *in vitro* (Devalla et al., 2015; Hochgreb, 2003; Lee et al., 2017; Xavier-Neto et al., 1999). As we identified the Notch pathway as being enriched in ventricular cardiac mesoderm, we hypothesized that activating Notch would increase ventricular cardiomyocyte specification. To assess how Notch activation in the CMes affects the phenotype of the differentiated cardiomyocytes, we FACS-purified cardiomyocytes at day 30 and performed RNA sequencing. Cardiomyocytes from Notch-induced CMes were distinct from those derived from both untreated (control) and DAPT-treated CMes cultures (Fig. 4a). PCA illustrates that PC1 captures the majority of the variance (67%) and largely separates cardiomyocytes from Notch-induced CMes from the other samples, while PC2 (18% variance captured) reflects experiment-to-experiment variability, even after batch correction with adjustment for hidden covariance. Differential gene expression confirmed the similarity between control and DAPT-treated cultures, yielding no differentially expressed transcripts (FDR<10%, fold change>1.5, base mean>2). However, cardiomyocytes from Notch-induced cultures express 1,554 transcripts differentially (443 upregulated, green points; 1,111 downregulated, gray points; FDR<10%, fold change>1.5) when compared to control cardiomyocytes (Fig. 4b).

**Figure 4:**
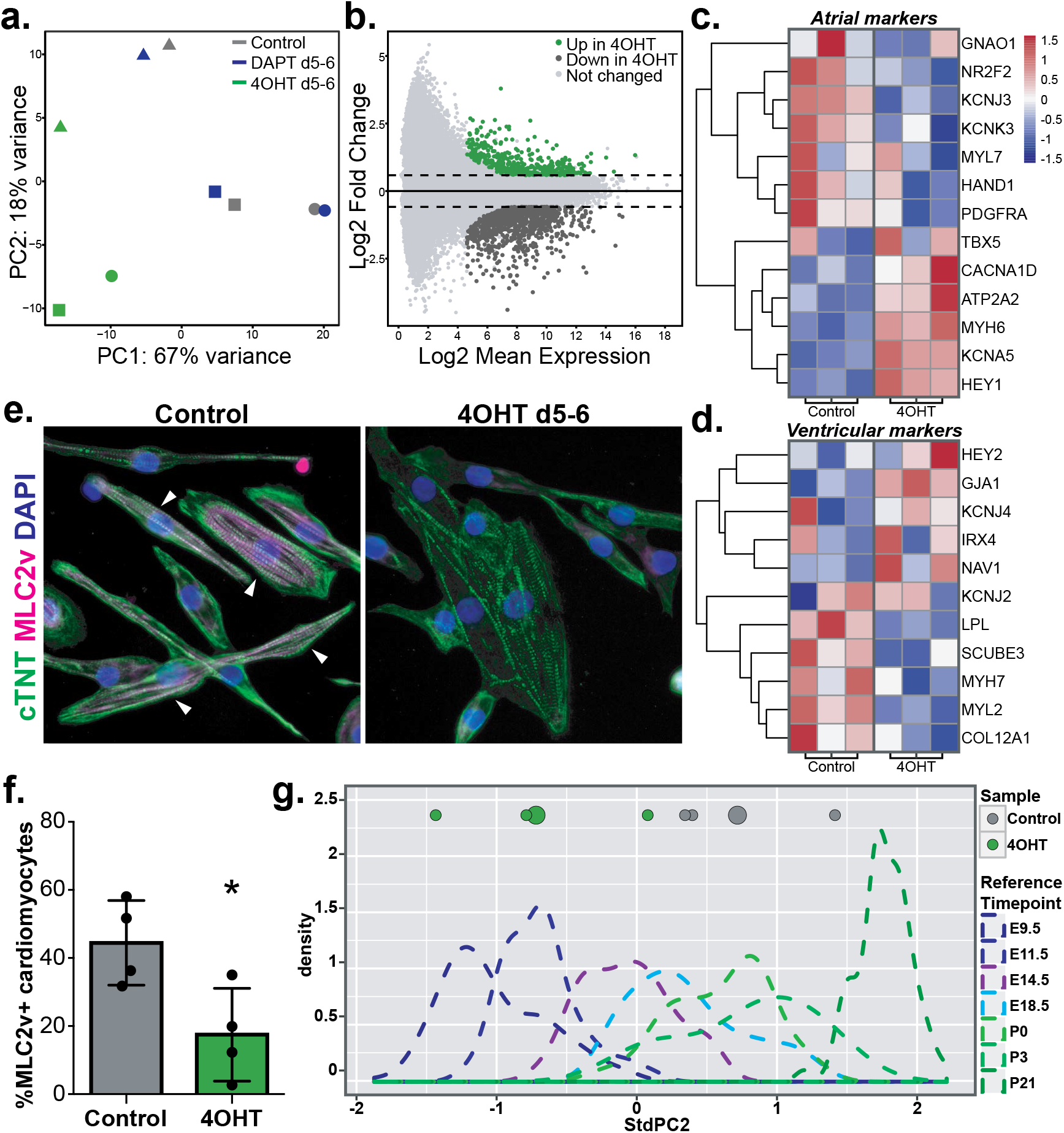
Gene expression analysis of cardiomyocytes derived from control or Notch-perturbed cardiac mesoderm. (a) Principal component analysis of RNA-sequencing data of cardiomyocytes from control, Notch inhibited (DAPT d5-6), or Notch induced (4OHT d5-6) cardiac mesoderm. Colors indicate cell type, symbols indicate replicates. (b) MA plot showing differentially expressed genes between cardiomyocytes from control or Notch induced cultures. (c-d) Gene expression z-scores for representative atrial (c) or ventricular (d) marker genes (genes taken from (Cyganek et al., 2018)).(e) Immunofluorescence analysis of cardiomyocytes from control or Notch induced cultures with antibodies against cTNT and the ventricular marker MLC2v. Arrowheads indicate MLC2v-expressing cardiomyocytes. (f) Flow cytometric quantitation of MLC2v-expressing cardiomyocytes from control or Notch induced cultures. (g) PCA-based determination of cardiomyocyte maturation. Reference stage data (dotted lines) is adapted from (DeLaughter et al., 2016). PC values of comparison data for individual replicates are shown as small circles and averages shown as large circles.

As Notch was enriched in vCMes *in vivo*, we analyzed whether Notch induction in hPSC-derived CMes biases differentiation toward chamber-specific fates. Expression of several well-established atrial and ventricular marker genes did not show a consistent trend, suggesting that Notch signaling does not act to induce a chamber-specific lineage specification mechanism at this early stage of development (Fig. 4c, d) (Cyganek et al., 2018). Interestingly, the expression of *MYL2* (MLC2v), a ventricular-specific gene expressed during late differentiation, is downregulated in cardiomyocytes from Notch-induced CMes cultures compared to control (Fig. 4c). We confirmed reduced MLC2v expression at the protein level through immunofluorescence analyses and quantification by flow cytometry (Fig. 4e, f). However, no difference was seen in *IRX4* expression, the earliest expressed ventricular-specific transcription factor, suggesting that while ventricular differentiation is compromised, fate acquisition may be unchanged (Fig. 4c).

As an alternative hypothesis, we explored whether the early induction of Notch signaling changes the maturation state of the resulting cardiomyocytes. *MYH6* and *MYH7*, which are often used as atrial and ventricular markers, respectively, are expressed dynamically during development. Specifically, *MYH6* is expressed throughout the early heart tube and downregulated as the ventricles develop, concomitant with *MYH7* upregulation (England and Loughna, 2013). Supporting our hypothesis, cardiomyocytes from Notch-induced CMes cultures expressed higher levels of *MYH6* and lower levels of *MYH7*, suggesting that they may be arrested in an immature state when compared to control cardiomyocytes (Fig. 4c, d). To analyze this in an unbiased fashion, we adapted a recently published principal component-based approach to assess the developmental stage of *in vitro*-derived cardiomyocytes (DeLaughter et al., 2016). This analysis leverages single cell RNA-seq data from mouse hearts collected from E9.5 to postnatal day (P) 21. By using a selection of genes expressed in a majority of P0 left ventricle cells, principal component analysis separates the data based on developmental time (Fig. 4g, dotted lines). We applied this same analysis to the gene expression data of cardiomyocytes from control and Notch-induced CMes cultures and overlaid the results onto the mouse scRNA-seq reference data to determine their relative developmental maturity. This analysis revealed that the cardiomyocytes from Notch-induced CMes cultures are more similar to cardiomyocytes from earlier developmental stages when compared to control cardiomyocytes (Fig. 4g), suggesting that Notch induction during early differentiation increases cardiomyocyte generation but leads to permanent changes that arrest the cells in an immature state. This finding is in conflict with recent data showing that treatment with polyinosinic-polycytidylic acid (pIC) resulted in an increase in Notch signaling and concomitant maturation of cardiomyocytes (Biermann et al., 2019). However, pIC treatment also inhibited TGFβ signaling in these experiments, so it is unclear whether Notch has differing effects depending on the broader signaling environment.

### Notch acts in a mesoderm-specific manner to enhance cardiomyocyte differentiation

We next investigated the mechanism by which active Notch signaling increases cardiomyocyte differentiation efficiency. Notch is a well-known regulator of proliferation, leading us to hypothesize that the observed increase in cardiomyocytes is due to an expansion of early cardiac precursor cells (Artavanis-Tsakonas et al., 1999). We therefore assessed proliferation over the course of differentiation and compared control and Notch-induced differentiation cultures (Sup. Fig. 5a). Cells were collected daily from day 5-10 and on days 15 and 30, after being cultured in the presence of the thymidine analog EdU for 24 hours prior to collection and analysis. Cells treated with 4OHT from days 5-6 showed a small increase in EdU incorporation at day 6 and 7, but this increase diminished quickly thereafter (Sup. Fig. 5b). We therefore conclude that proliferation is not the key mechanism by which enhanced Notch signaling increases cardiomyocyte differentiation efficiency.

An alternative hypothesis is that Notch induction causes conversion of non-cardiac progenitors to the cardiac lineage. Since little is known about the role of Notch signaling within gastrulation-stage populations, we sought to use the hPSC differentiation system to dissect the mechanism of its action on these cells. Mesoderm can be detected as early as day 4 of differentiation using the cell surface marker CD13, and CXCR4 is used to exclude endoderm from the analysis (Skelton et al., 2016). We first asked whether Notch increases the commitment of mesoderm to the cardiomyocyte fate or whether it can also convert non-mesodermal cells to the cardiac lineage. We FACS purified mesoderm (CD13+CXCR4-) and non-mesoderm (CD13-) populations at day 4 and plated them as individual monolayers for continued differentiation. At day 5, a subset was treated with 4OHT to induce Notch signaling from day 5-6, as done previously. At day 30 the cultures were analyzed via flow cytometry to quantify differentiation efficiency (Fig. 5a). Differentiation cultures at day 4 were composed of 60±4.7% mesoderm and 20±5% non-mesodermal cells (Fig. 5b). Mesoderm differentiated in the absence of Notch induction yielded 30±2.5% cardiomyocytes at day 30 (Fig. 5c, left). However, when Notch was induced from day 5-6, the efficiency of differentiation was increased to 80±0.7% (Fig. 5c, right). As expected, non-mesoderm cells yielded very few cardiomyocytes at day 30. Induction of Notch signaling did not result in an increase when compared to control cultures, indicating that non-mesoderm cells are not converted to the cardiac lineage upon Notch activation (Fig. 5d).

**Figure 5:**
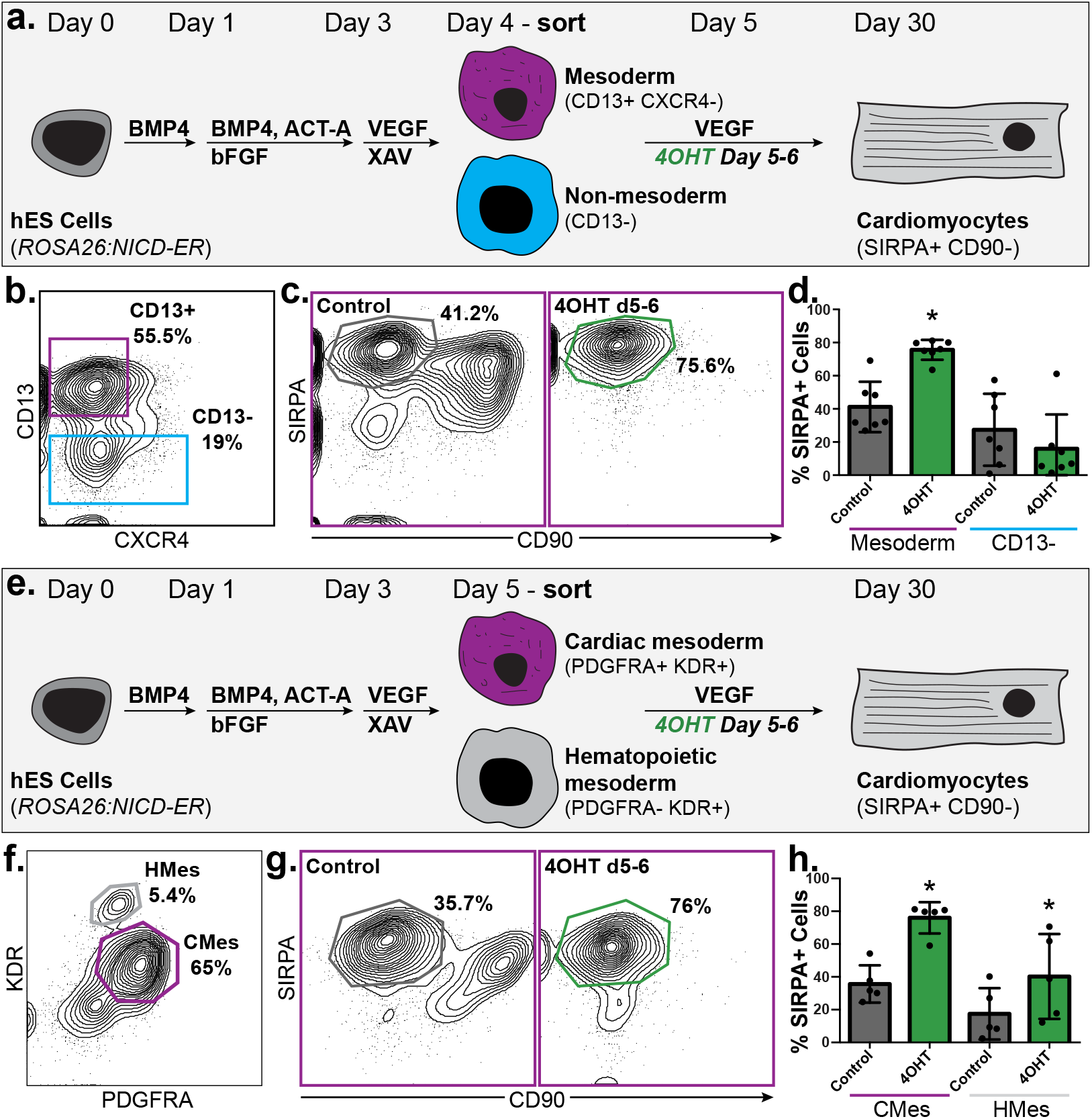
Notch acts in a mesoderm-specific manner to enhance cardiomyocyte differentiation. (a) Schematic of *RO-SA26:NICD-ER* hPSC-derived day 4 mesoderm purification and cardiomyocyte differentiation. (b) FACS isolation to purify CD13+CXCR4-mesoderm and CD13-non-mesoderm on day 4 of differentiation. (c) Flow cytometric analysis of cardiomyocyte generation from CD13+CX-CR4-mesoderm in the absence or presence of Notch induction from day 5-6. (d) Quantification of cardiomyocyte differentiation from mesoderm and non-mesoderm. (e) Schematic of *RO-SA26:NICD-ER* hPSC-derived day 5 mesoderm purification and cardiomyocyte differentiation. (f) FACS isolation to purify PDG-FRA+KDR+ cardiac mesoderm and PDGFRA-KDR+ hematopoietic mesoderm. (g) Flow cytometric analysis of cardiomyocyte generation from PDGFRA+K-DR+ cardiac mesoderm in the absence or presence of Notch induction from day 5-6. (h) Quantification of cardiomyocyte differentiation efficiency from cardiac mesoderm and hematopoietic mesoderm.

To confirm the mesoderm-specific effect seen in these experiments, we differentiated hPSCs into definitive endoderm progenitors, induced Notch, and analyzed for the presence of cardiomyocytes to assess fate conversion (Sup. Fig. 6a). Definitive endoderm was generated through the addition of high amounts of Activin A at days 1 and 4 of differentiation, and cells were treated with 4OHT from day 5-6, as in previous experiments. This approach yields 49.5±23.6% definitive endoderm cells at day 5 of differentiation, as indicated by the expression of CXCR4 and cKIT (Sup. Fig. 6b) (D’Amour et al., 2005; Gouon-Evans et al., 2006). A very small number of cardiomyocytes (5.1±2% SIRPA+ CD90-) were observed at day 30, and Notch induction was not effective at specifying additional cardiomyocytes (Sup. Fig. 6c, d). Thus, consistent with the above results, Notch pathway activity has no cardiogenic effect in non-mesoderm cell types, and the increase in cardiomyocyte differentiation observed is due to a mesoderm-specific mechanism.

Our data thus far show that Notch acts in a mesoderm-specific fashion to increase cardiomyocyte differentiation efficiency. We next determined if the effects of Notch signaling are restricted to the cardiac lineage or can convert other mesoderm lineages to the cardiac fate. Other mesoderm progenitors are generated during differentiation of hPSCs to cardiomyocytes and can be identified by specific cell surface markers. At day 5 of differentiation, cardiac mesoderm can be identified via surface expression of PDGFRA and KDR, consistent with data described the mouse system (Kataoka et al., 1997; Kattman et al., 2006; Kattman et al., 2011; Takakura et al., 1997). Hematopoietic mesoderm is also generated and characterized by high expression of KDR (KDR^hi^). It has been previously shown that mouse PSC-derived hematopoietic mesoderm can be differentiated to cardiomyocytes through Notch induction (Chen et al., 2008). We therefore wanted to determine if Notch signaling is sufficient to induce cardiomyocyte differentiation from hematopoietic mesoderm in human cells as well. We FACS purified CMes (PDGFRA+KDR+) and HMes (KDR^hi^) at day 5 and plated each as monolayers for continued differentiation. A subset was treated with 4OHT to induce Notch signaling at the time of plating, and cardiomyocyte generation was measured at day 30 (Fig. 5e). Day 5 cultures comprised 53.8±17% CMes and 4.7±1.1% HMes (Fig. 5f). CMes that was differentiated without Notch induction gave rise to 35.7±11.4% SIR-PA+CD90-cardiomyocytes, similar to the control CD13+CDCR4-day 4 mesoderm (Fig. 5g, left). Again, Notch induction increased this efficiency to 76±9.5% (Fig. 5g, right). We also observed a statistically significant increase when differentiating HMes into cardiomyocytes in the presence of Notch signaling as compared to control, consistent with previous data from the mouse PSC system (Fig. 5h). Overall, our data from these experiments suggest that germ layer fate switching is not a major contributing factor to the observed increase in differentiation efficiency in Notch induced cultures when compared to controls. Rather, these data support the hypothesis that Notch acts to enhance commitment of the mesoderm progenitors to the cardiac lineage.

The segregation of atrial and ventricular lineages occurs very early during cardiomyocyte differentiation (Bardot et al., 2017b; Devalla et al., 2015; Devine et al., 2014; Lee et al., 2017; Lescroart et al., 2014). All of the experiments to this point have been performed with a protocol that yields primarily ventricular cardiomyocytes. To determine if Notch signaling has similar effects on atrial and ventricular cardiac mesoderm, we next assessed the effect of Notch signaling on atrial cardiomyocyte differentiation. Atrial cardiomyocytes can be differentiated through the addition of all-trans retinoic acid (atRA) at day 3 of differentiation (Sup. Fig. 6e) (Devalla et al., 2015; Lee et al., 2017). In addition to generating atrial-like cardiomyocytes, adding atRA also greatly increases the efficiency of differentiation (Sup. Fig. 6f, g). We did not observe a further increase in cardiomyocyte differentiation efficiency upon Notch induction from day 5-6 (Sup. Fig. 6f, g).

### Notch induction is required for cardiomyocyte differentiation in the absence of WNT inhibition

Given that Notch signaling appears to act on mesoderm specifically, we sought to determine whether Notch interacts with other signaling pathways during cardiac commitment. At the time points when Notch-induction was most effective (day 4-5 or 5-6), cells are cultured in the presence of VEGF and the WNT inhibitor XAV939 (days 3 to 5), or VEGF alone (days 5 to 7), suggesting that VEGF or WNT signaling may play parallel or additive roles with Notch at this time. It has been shown *in vivo* that there is a switch from canonical to non-canonical Wnt signaling at this stage, and that Notch may play a role in this process (Guo et al., 2019; Klaus et al., 2012; Miazga and McLaughlin, 2009; Wang et al., 2018). To test whether Notch signaling is sufficient to specify mesoderm to the cardiac lineage, we first excluded VEGF during differentiation, and added 4OHT to induce Notch signaling from day 4-5 (Sup. Fig. 7a). Culturing cells in the absence of VEGF throughout differentiation had no effect on overall cardiomyocyte generation efficiency, whether or not Notch was induced (Sup. Fig. 7b, c). Immunofluorescence analysis for cTNT and alpha-actinin (aACT) confirmed that phenotypically normal cardiomyocytes were present in the absence of VEGF (Sup. Fig. 7d), leading us to conclude that under these culture conditions VEGF has no effect on hPSC differentiation to cardiomyocytes.

However, cells differentiated without XAV939 yielded less than 8.5±4% cardiomyocytes at day 30, confirming that WNT inhibition is essential for commitment of cells to the cardiac lineage, as shown previously (Fig. 6a, b) (Lian et al., 2013). Addition of DAPT failed to enhance this effect (Fig. 6b). Interestingly, induction of Notch from day 4-5 in the absence of XAV939 was able to restore cardiomyocyte differentiation to control levels (Fig. 6b). This suggests that Notch is required for cardiomyocyte differentiation in the absence of WNT inhibition, and that Notch may be acting to inhibit WNT signaling during that process. To further examine that hypothesis, we cultured cells in the presence of the GSK3B inhibitor CHIR, which stabilizes B-catenin and results in constitutive activation of canonical WNT signaling. Addition of CHIR completely blocked cardiomyocyte differentiation, and Notch induction failed to rescue this effect (Fig. 6b). Quantification of these data confirm the interaction of Notch and WNT signaling in these experiments (Fig. 6c). In all conditions where cardiomyocytes were detected by flow cytometry analysis, immunofluorescence analysis for cTNT and aACT confirmed the presence of phenotypically normal cardiomyocytes (Fig. 6d). Overall, our data support a model in which Notch signaling acts to increase cardiac lineage commitment by inhibiting WNT activity and show that Notch signaling on its own is sufficient to commit mesoderm to the cardiac lineage.

**Figure 6:**
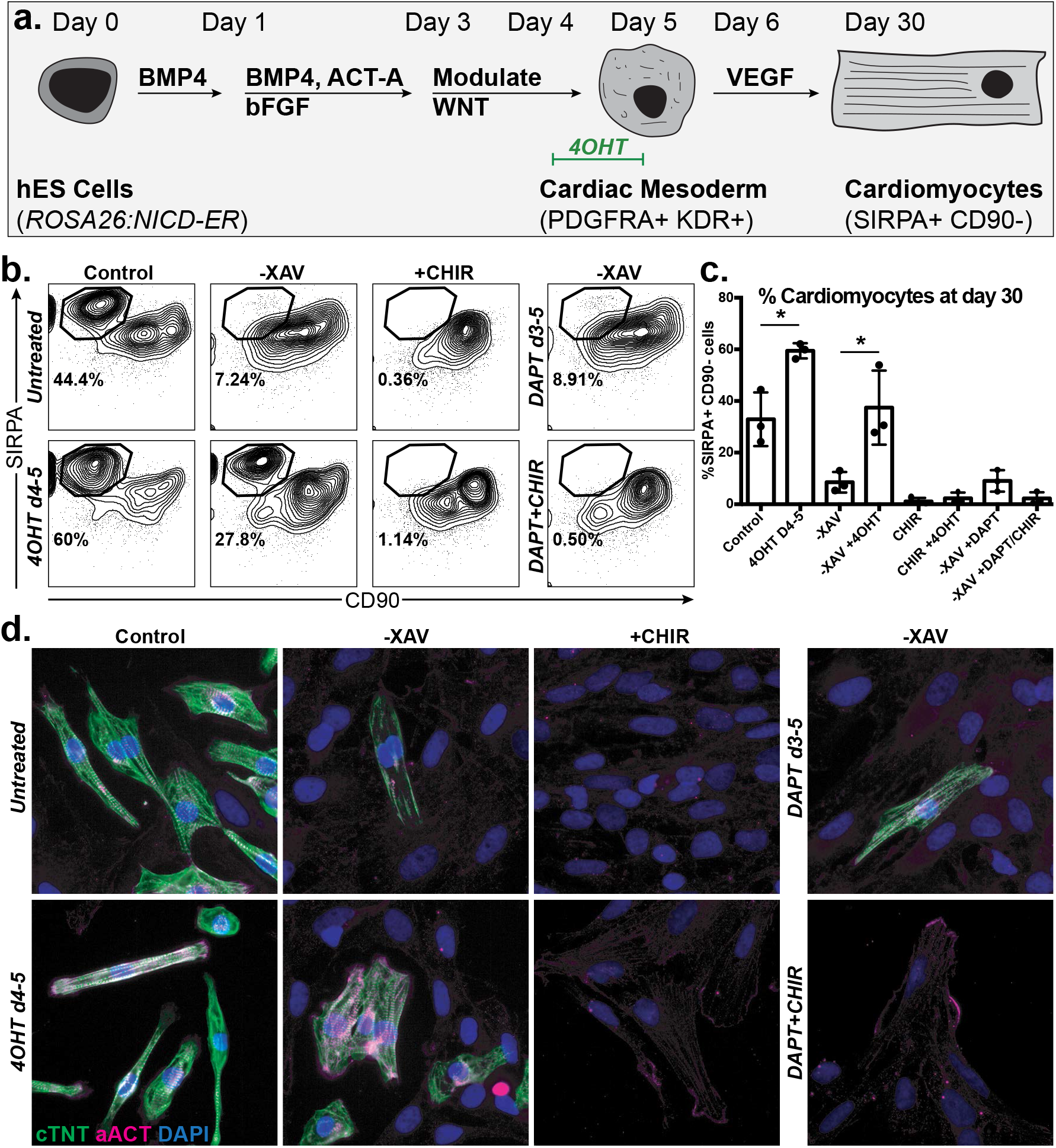
Notch induction is required for cardiomyocyte differentiation in the absence of WNT inhibition. (a) Schematic of *ROSA26:NICD-ER* hPSC-cardiomyocyte differentiation with WNT and Notch modulation. (b) Flow cytometric analysis of cardiomyocyte generation in the absence of WNT inhibition (-XAV) or presence of WNT induction (CHIR), with and without Notch induction (4OHT) or inhibition (DAPT). (c) Quantification of data in (b). (d) Immunofluorescence analysis on day 30 cardiomyocytes with antibodies against cTNT and aACT.

## Discussion

We have previously identified a ventricular-specific cardiac progenitor population specified during gastrulation through the use of *Foxa2* lineage-tracing (Bardot et al., 2017b). Here, we have expanded on these findings and studied the heterogeneity of cardiac mesoderm cell populations and the early steps of cardiac mesoderm differentiation both *in vivo* and *in vitro*. Our study focused on a developmental stage occurring at E7.25 *in vivo* or day 5 of hPSC differentiation, which is likely prior to lineage commitment. This stage encompasses the earliest steps of cardiac specification and captures the differential embryonic signaling environment cells are exposed to as they migrate anteriorly to form the cardiac crescent. *Foxa2* lineage-tracing has enabled us to distinguish, at this early stage, the mesoderm populations that will give rise to ventricular cells (vCMes) versus atrial cells (aCMes) later during development, and therefore gives us the ability to investigate early versus late differentiating cells of the cardiovascular lineage.

The *in vivo* RNA-seq analysis we performed suggest that cardiac mesoderm from the mouse embryo and from mESCs are overall similar after accounting for technical differences. Notch signaling appears to be active in all cardiac mesoderm populations, though cells further along the differentiation trajectory in both systems (Foxa2-hCD4+ CMes *in vitro*, Foxa2Cre:YFP+ CMes *in vivo*) express higher levels of some Notch pathway components. Other studies have shown that *Notch1* is expressed in early Mesp1 + cardiac precursors that are fated for the endothelial lineage, but did not address whether Notch signaling plays a role in that lineage decision (Lescroart et al., 2018). We observed expression of *Notch1-3* in both aCMes and vCMes, though we are unable to determine the relationship between receptor expression and lineage decision from our data. By taking advantage of the hPSC cardiomyocyte differentiation model, we were able to show that inducing Notch signaling in hPSC-derived cardiac mesoderm enhances cardiomyocyte differentiation overall. Such transient Notch induction appears not to affect atrial vs. ventricular fate decisions, but rather increases cardiomyocyte generation efficiency more generally. This suggests that while Notch components are enriched in ventricular-fated cardiac mesoderm, we likely capture a snapshot of development when Notch signaling is being consecutively activated in migratory mesoderm. While all cardiac mesoderm will be subjected to Notch signaling, the early differentiating ventricular progenitors receive the signal first, while later-specified atrial progenitors are exposed to it subsequently.

Within the heart, Notch is known to act at numerous stages during development in different organisms (High and Epstein, 2008). Aberrant activation of Notch signaling in cardiac precursors causes an enlargement of the left ventricle and may negatively impact maturation of the forming cardiomyocytes (Watanabe, 2006). While Notch activation in the cardiac lineage showed a hyper-plasic phenotype, induction of Notch signaling in the epiblast has a negative overall impact on mesoderm specification and organization (Souilhol et al., 2015). Data from Xenopus suggests that active Notch signaling is important during a brief time window between gastrulation and differentiation, and that this timing acts to coordinate progenitor maintenance and differentiation. Specifically, premature dampening of the Notch pathway results in cardiac progenitors upregulating the differentiation program prior to reaching the midline of the embryo (Miazga and McLaughlin, 2009). This is consistent with our data showing that transient Notch induction specifically at the cardiac mesoderm stage enhances cardiomyocyte differentiation efficiency. We also showed that the inductive effects of Notch signaling are specific to mesoderm, and that Notch was insufficient to convert endoderm to the cardiac lineage.

Knowing that the differentiation protocol used provides a highly cardiogenic signaling environment, we sought to determine the interaction between Notch and other relevant pathways. Despite the expression of *VEGFR2* (KDR) in both *in vivo* and *in vitro* mesoderm, removing VEGF had little effect on hPSC differentiation overall. This is consistent with data from the mouse embryo showing that while *Kdr* is required for the formation of hematopoietic lineages, it is not required for cardiac progenitor formation (Ema et al., 2006). Notch is known to interact with other signaling pathways, particularly the BMP and Wnt pathways. Notch has been shown to activate the non-canonical *Wnt5a* ligand later during heart development, further suggesting a link between these two pathways (Wang et al., 2018). It was previously shown that mouse ESC-derived Kdr+ cells fated to the hematopoietic and vascular lineages can be redirected to the cardiac lineage via Notch activation (Chen et al., 2008). Gene expression profiling revealed that this was in part due to upregulation of BMP pathway genes and downregulation of canonical Wnt signaling, ultimately mimicking the signaling cocktail used to generate cardiomyocytes from PSCs. In the embryo, the overlying endoderm provides high levels of Wnt-inhibitory proteins to the developing mesoderm, and proper endoderm development has been shown to be critical for heart development (Marvin et al., 2001; Nascone and Mercola, 1995; Schneider and Mercola, 2001; Schultheiss et al., 1995).

In accordance with an important role for Wnt inhibition during cardiac mesoderm specification, our analysis reveals expression of secreted Wnt inhibitory proteins and non-canonical ligands, with *Sfrp1* being enriched in early-migrating ventricular cardiac mesoderm (Figure 2b). Consistent with the importance of that interaction, we found that failing to inhibit WNT *in vitro* almost completely eliminated cardiomyocyte differentiation. Interestingly, Notch induction is sufficient to rescue hPSC differentiation to control levels, providing evidence that these two pathways interact during cardiac mesoderm commitment. We therefore believe that Notch in the cardiac mesoderm is acting in concert with the overlying endoderm to establish a Wnt-inhibitory zone as cells approach the anterior side of the embryo, and that inducing Notch during hPSC differentiation recapitulates aspects of this mechanism. These findings have implications for therapeutic uses, as a fuller understanding of the *in vivo* signaling environment may lead to development of cardiomyocyte differentiation protocols with improved efficiency and cardiomyocyte subtype selectivity.

Our experiments illustrating enhanced cardiomyocyte differentiation after Notch induction nicely corroborate our gene expression profiling *in vivo* and suggest that Notch plays a role during early mammalian heart development, a stage that is technically challenging to manipulate in a temporally and spatially controlled manner. Going forward, it will be important to dissect the detailed mechanism for how this occurs, and whether Notch signaling in the cardiac mesoderm is broadly cardiogenic. While studies have looked at the effects of Notch signaling on mouse development, these have largely been done using constitutive Notch activation (Souilhol et al., 2015; Watanabe, 2006). It would be of great interest to perturb Notch signaling within the early cardiac lineage in a temporally controlled and reversible manner, as has been done in other model organisms (Miazga and McLaughlin, 2009).

In summary, we have used a combination of mouse embryo and human PSC approaches to dissect the early steps of heart progenitor specification. By performing whole transcriptome analysis on *in vivo* cardiac mesoderm, we identified an enrichment for Notch pathway components in the early-differentiating ventricular lineage. Using human pluripotent stem cell directed differentiations, we found that transient Notch induction at the cardiac mesoderm stage increases the efficiency of cardiomyocyte differentiation. Surprisingly, Notch activation does not impact the proportion of ventricular cardiomyocytes formed, but rather yields cells less developmentally mature than control counterparts. Finally, we show that the cardiac-inducing effect of Notch signaling is restricted to mesodermal cells and may act to inhibit Wnt signaling. These data reveal a role for Notch signaling in committing early mesoderm to the cardiac lineage in both mouse and human systems.

## Materials and methods

### Mice

The *Foxa2Cre* mouse line (C57BL/6J) was generated and shared with us by Dr. Heiko Lickert (Horn et al., 2012). *Rosa26-YFP* (006148; C57BL/6J), *Rosa26-mTmG* (007676; C57BL/6J), and *Rosa26-H2B-mCherry* (023139; B6/129S) mice were all obtained from The Jackson Laboratory. Studies were performed on mixed backgrounds as dictated by strain availability. For time-course experiments, the day of plug identification corresponds to embryonic day 0.5 (E0.5). All animals were housed in facilities operated by the Center for Comparative Medicine and Surgery (CCMS) at Icahn School of Medicine. All animal experiments were conducted in accordance with the guidelines and approval of the Institutional Animal Care and Use Committee at Icahn School of Medicine at Mount Sinai.

### Cell lines

The *ROSA26:NICD-ER* hESC line was generated and generously shared with us by Dr. Paul Gadue (Ditadi et al., 2015; Gadue et al., 2006). The *MYL2-mEGFP* WT hiPSC line was obtained from the Coriell Institute (Line no. AICS-0060-027).

### Histology and Immunofluorescence

Embryos were collected from timed pregnancies and either fixed in 4% PFA, washed with phosphate buffered saline (PBS) and equilibrated in 30% sucrose (Sigma-Aldrich) before embedding, or embedded fresh into OCT (Electron Microscopy Sciences). Tissues were subsequently cut into 10μm sections using a Leica Cryostat. Slides were fixed for 10 min in 4% PFA and incubated for 1 h in blocking solution (PBS with 0.1%Triton and 1% BSA). Primary antibodies were diluted in blocking solution and incubations were carried out for 1 h at room temperature or overnight at 4°C, followed by incubation in secondary antibody for 1 h at room temperature. Slides were then counterstained with DAPI and mounted using glycerol-based nPG (Sigma-Aldrich, P3130) antifade mounting media.

For whole mount immunofluorescence, embryos were collected, immersion-fixed in 4% PFA overnight, and washed with PBS. Embryos were blocked for >4 h in blocking solution (PBS with 0.1%Triton and 1% BSA). Primary antibodies were diluted in blocking solution and incubations were carried out overnight at 4°C. Embryos were washed three times in PBS-T (PBS with 0.1%Triton) for 2 h each, followed by incubation in secondary antibody overnight at 4°C. Embryos were then counterstained with DAPI and allowed to equilibrate in nPG antifade mounting media before mounting. Embryos were mounted on slides using double-stick tape and coverslips. The following primary antibodies and dilutions were used: anti-GFP (abcam, 1:500); anti-RFP (Rockland, 1:500); anti-cTnT (ThermoSci, 1:300); anti-Mlc2v (Proteintech, 1:100). Secondary antibodies conjugated with Alexa dyes were obtained from Jackson Immunoresearch and used at 1:500.

Fluorescence images were obtained using either Leica DM6000 or DM5500B microscopes. Confocal microscopy was performed using a Zeiss 780 microscope. Images were processed using Imaris 8, FIJI ImageJ, or Adobe Photoshop software (Bardot et al., 2017a).

### Human pluripotent stem cell differentiations

Mesoderm/Cardiac: Embryoid bodies (EBs) were generated from ESC lines using collagenase B (Roche) and cultured in RPMI 1640 base media (ThermoFisher) containing 0.5X B27 supplement (ThermoFisher), 2mM glutamine (Gibco-BRL), 1 mM ascorbic acid (Sigma), and 4×10-4 M monothioglycerol (Sigma)) with BMP4 (1 ng/ml, R&Dsystem) and Thiazovivin (2 μM, Millipore). On day 1, EBs were harvested and resuspended in induction medium (base media with basic fibroblast growth factor (bFGF; 2.5 ng/ml, R&Dsystem), activin A (20 ng/ml, R&Dsystem) and BMP4 (20 ng/ ml)). On day 3, the EBs were harvested and resuspended in base media with vascular endothelial growth factor (VEGF; 10 ng/ml, R&Dsystem) and XAV939 (10 μM, Stemolecule). On day 3, the EBs were harvested and resuspended in base media with vascular endothelial growth factor (VEGF; 10 ng/ml, R&Dsystem). On day 8 and thereafter, EBs were cultured in base media until day 30 for further analysis.

Endoderm: Human pluripotent stem cells were differentiated into the endoderm lineage as previously described (Goldman et al., 2013). Briefly, cells were dissociated using collagenase B (Roche) and cultured in low cluster plates to allow EB formation in in RPMI 1640 base media (ThermoFisher) supplemented with BMP4 (3 ng/ml, R&D Systems) and Thiazovivin (2 μM, Millipore). At day 1 the medium was changed to base media supplemented with Activin A (100 ng/ml, R&D Systems), basic FGF (bFGF, 2.5 ng/ml, R&D Systems), and BMP4 (0.5 ng/ml, R&D Systems). At day 4, the medium was changed to the base media supplemented with Activin A (100 ng/ml), bFGF (2.5 ng/ml), and vascular endothelial growth factor (VEGF, 10 ng/ ml, R&D Systems). On day 5 EBs were dissociated and analyzed for CXCR4+ cKIT+ cells. On day 6 and thereafter, EBs were cultured in base media until day 30 for further analysis.

### Isolation and differentiation of early subpopulations during differentiation

On day 4 of cardiac differentiation, EBs were dissociated using TrypLE (Invitrogen). CXCR4-CD13+ and CXCR4-CD13+ cells were sorted and cultured in matrigel-coated 96 well plates at 1 × 105 cells per well in base media with vascular endothelial growth factor (VEGF; 10 ng/ml), XAV939 (10microM), and Thiazovivin (2 μM). On day 5, the media was changed to base media with vascular endothelial growth factor (VEGF; 10 ng/ml), and cultures were maintained in this media until day 8 of differentiation, when cells were cultured in base media only, with media changes every 3 days until day 30, when cells were analyzed.

On day 5 of cardiac differentiation, EBs were dissociated using TrypLE (Invitrogen). PDGFRA+K-DR+, PDGFRA-KDR+ and PDGFRA-KDR-cells were sorted and cultured in matrigel-coated 96 well plates at 1×105 cells per well in base media with vascular endothelial growth factor (VEGF; 10 ng/ml) and Thiazovivin (2 μM). On day 6, the media was changed to base media with vascular endothelial growth factor (VEGF; 10 ng/ml), and cultures were maintained in this media until day 8 of differentiation, when cells were cultured in base media only, with media changes every 3 days until day 30, when cells were analyzed.

### Flow cytometry analysis and cell sorting

Mouse embryos: E7.25 embryos were dissociated with TrypLE. Cells were washed in staining solution (DMEM with 0.1% BSA), pelleted, and resuspended in staining solution. Antibodies were diluted in staining solution and cells were incubated on ice for 30 min. Cells were then washed, filtered, and resuspended in staining solution for analysis or cell sorting.

Day 3 to day 15 hPSC-EBs: EBs generated from hPSC differentiation experiments were dissociated with 0.25% trypsin/EDTA.

Day 20 and older hPSC-EBs: EBs were incubated in collagenase type II (1 mg/ml; Worthington) in Hanks solution (NaCl, 136 Mm; NaHCO3, 4.16 mM; NaPO4, 0.34 mM; KCl, 5.36 mM; KH2PO4, 0.44 mM; dextrose, 5.55 mM; HEPES, 5 mM) overnight at 37 °C. The EBs were pipetted gently to dissociate the cells, washed, and resuspended in staining solution (PBS with 0.1% BSA) and then filtered. Antibodies were diluted in staining solution and cells were incubated on ice for 30 min. Cells were then washed and resuspended in staining solution for analysis or cell sorting.

Cell counts were collected using a LSRII (BD Biosciences) and data was analyzed using the FlowJo software.

The following antibodies and dilutions were used: anti-mouse Pdgfra-BV421 (BD, 1:100); anti-mouse Kdr-PE-Cy7 (BD, 1:100); anti-mouse

Epcam (eBioscience, 1:100); anti-cardiac Troponin T (ThermoSci, 1:100); anti-human CD172a/b (SIRPα/β) (Biolegend; clone SE5A5; 1:200) anti-human CD13 (Biolegend; clone WM15; 1:1,000), anti-human CD90 (Biolegend, clone 5E10; 1:200); anti-human CD140a (PDGFRα) (Biolegend, clone 16A1; 1:250); anti-human CD309 (KDR/VEGFR2) (Biolegend, clone 7D4-6; 1:200); anti-human CD184 (CXCR4) (Biolegend, clone 12G5; 1:500).

### cDNA library preparation and RNA-sequencing

Total RNA obtained from FACS-sorted E7.5 mouse embryonic populations was purified with the Absolutely RNA nanoprep kit (Agilent) and quantified as above. Sample cDNA libraries were prepared using the Ovation RNA-Seq System V2 (NuGEN). First strand cDNA was analyzed by qPCR, and only samples with faithful expression of the markers used for FACS isolation were submitted for sequencing.

Total RNA from FACS-sorted hPSC-derived cardiomyocytes was isolated using the Quick-RNA microprep kit (Zymo Research) and submitted to the NYU Genomics Facility for QC, library generation, and sequencing.

### RNA-seq data processing

Quality analysis of raw sequencing reads was performed using FastQC (http://www.bioinformatics.bbsrc.ac.uk/proiects/fastqc/). Reads were then aligned to the mouse reference genome (mm10) for mouse samples or the human reference genome (hg19) for human samples using STAR with the two-pass setting (Dobin et al., 2013), and a database of known splice iunctions from the ENCODE mm10/GRCm38 annotation for the initial alignment (http://www.gencodegenes.org/mousereleases/2.html). Picard tools (http://broadinstitute.github.io/picard/) was used to index and remove duplicate reads from the resulting SAM files, generating 16-25 million unique aligned reads per sample. From the resulting alignments, we used HTSeq (Anders et al., 2015) to generate read counts per transcript, and normalized these and performed differential expression analysis using DESeq2 (Love et al., 2014). Gene ontology analysis was performed using Panther GO (http://geneontology.org/page/go-enrichment-analysis). All heatmaps were generated using the R package pheatmap (v 1.0.8). PCA and MA plots were generated using the R packages ggplot2 (v 2.0.0) and DESeq2 (v 1.10.1). The developmental benchmarking was performed using the R package princomp on Log10 FPKM data, as previously described (DeLaughter et al., 2016).

### Statistical analysis

Statistical analysis was performed using Student’s t-test and is presented as statistically significant (*) with a p-value cutoff of p < 0.05. Unless specified, error bars indicate mean ± standard deviation.

## Acknowledgements

We would like to thank Dr. Paul Gadue for providing the *ROSA26-ICN-ER* hESC line and the *Foxa2-hCD4* mESC line. The *Foxa2Cre* mice were generated and generously shared by Dr. Heiko Lickert. We greatly appreciate Drs. Dan DeLaughter and Christine Seidman for their assistance with the cardiomyocyte maturity analysis. We would like to acknowledge the ISMMS shared resource facilities, in particular the microscopy (Drs. Deena Benson and Nikos Tzavaras), medical illustration (Ni-ka Ford), flow cytometry (Christopher Bare and Venu Pothula), and stem cell (Dr. Sunita D’Souza) core facilities. Dr. Ariana Heguy and the NYU Genome Technology Center provided guidance and expertise on RNA-sequencing sample preparation and performed all RNA-sequencing.

## Author contributions

ESB and NCD designed and performed experiments and analyzed the data. BJ and AJS analyzed the RNA-sequencing data. NW performed experiments and analyzed the data. AR and MR developed adapted protocols for generating RNA-sequencing libraries from *in vivo* samples. ESB and NCD wrote the manuscript with input from all authors.

## Data availability

The authors declare that all data supporting the findings of this study are available within the article and its supplementary information files, or from the corresponding author upon request. The RNA sequencing raw reads and deSeq2 data have been deposited in the NCBI GEO database under the accession codes GSE143226 and GSE143227.

---

**Supplementary Figure 1:**
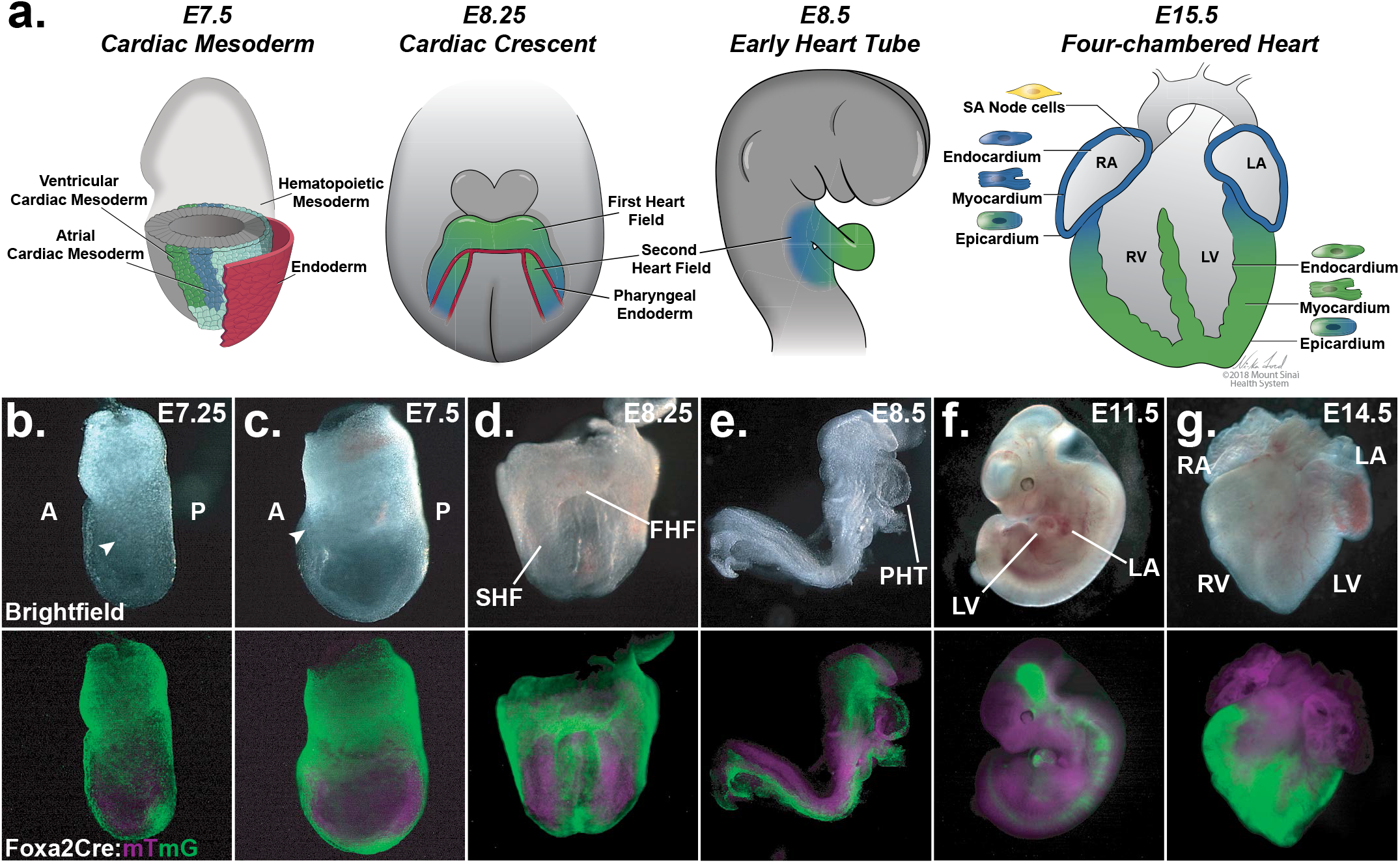
Distribution of Foxa2Cre:YFP lineage-traced cells during mouse development. (a) Schematic overview of ventricular progenitors during heart development. (b-g) Whole mount live imaging of *Foxa2Cre:mTmG* lineage tracing embryos. Early (b) and late (c) gastrulation stages show GFP+ migratory mesoderm. Cardiac crescent (d) and early heart tube (e) stage embryos show restriction of GFP+ cells to the prospective ventricular regions. GFP+ cells are restricted to the ventricles in E11.5 (f) and E14.5 (g) hearts.

**Supplementary Figure 2:**
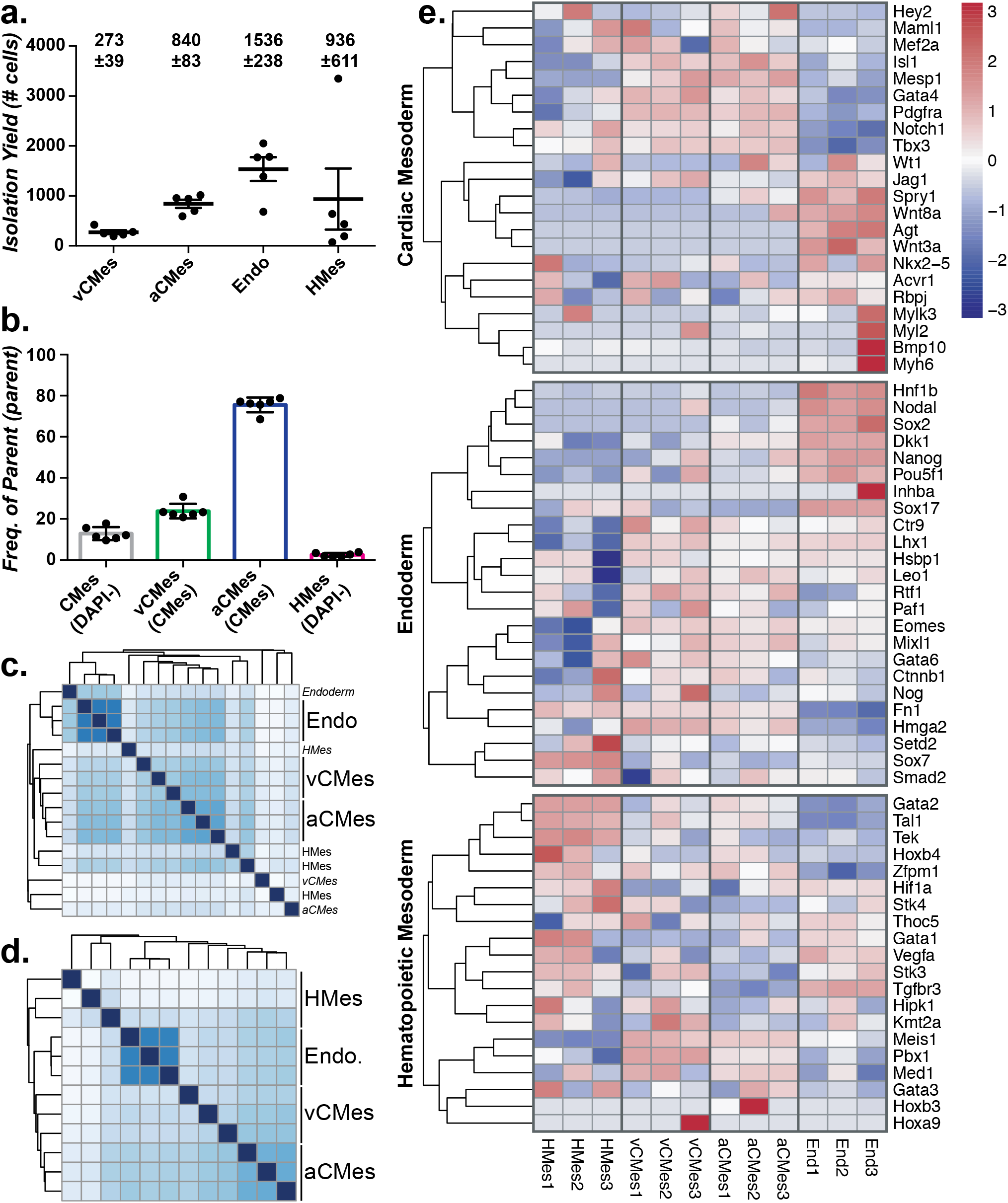
Isolation and gene expression profiling of embryonic progenitor populations. (a) Cell yield from FACS isolation experiments. E7.5 mouse embryos were stage-matched by morphology and pooled by litter. Cell numbers are for pools of 7-10 embryos. (b) Cardiac mesoderm contributed to approximately 10% of total viable cells. Ventricular cardiac mesoderm made up a smaller portion of total cardiac mesoderm than non-ventricular cardiac mesoderm. Hematopoietic mesoderm made up ~2.5% of the total viable population. (c, d) A total of four replicates were subjected to RNA sequencing and analyzed by hierarchical clustering. Based on this analysis, one replicate was removed prior to further analysis. (e) Expression z-scores for cardiac mesoderm, endoderm, and hematopoietic mesoderm signature genes.

**Supplementary Figure 3:**
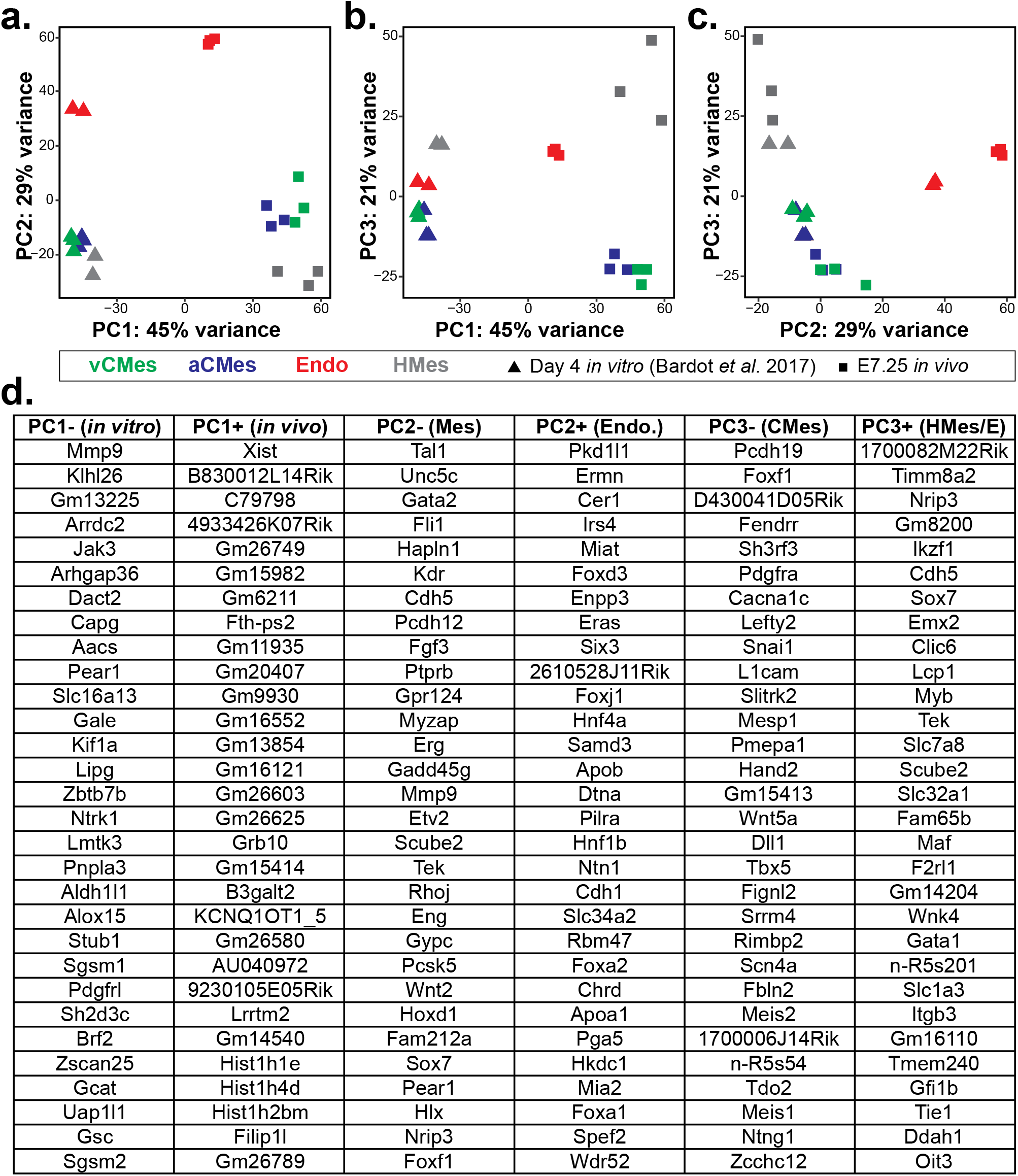
Comparison of RNA sequencing data from *in vitro* and *in vivo* populations of interest. (a) Principal component analysis of *in vitro* and *in vivo* cardiac mesoderm, endoderm, and hematopoietic mesoderm RNA sequencing data. The first three PCs capture 95% of the total variance across all samples. (a, b) PC1 separates samples based on cell origin and technical differences (*in vitro* vs. *in vivo*). (a) PC2 separates endoderm from mesodermal cell types, whereas (b) PC3 separates cardiac mesoderm from non-cardiac samples. (c) Comparison of PC2 vs. PC3 restores the lineage relationships observed when analyzing within *in vitro* or *in vivo* datasets. (d) Ordered gene lists for genes that explain variance within each principal component.

**Supplementary Figure 4:**
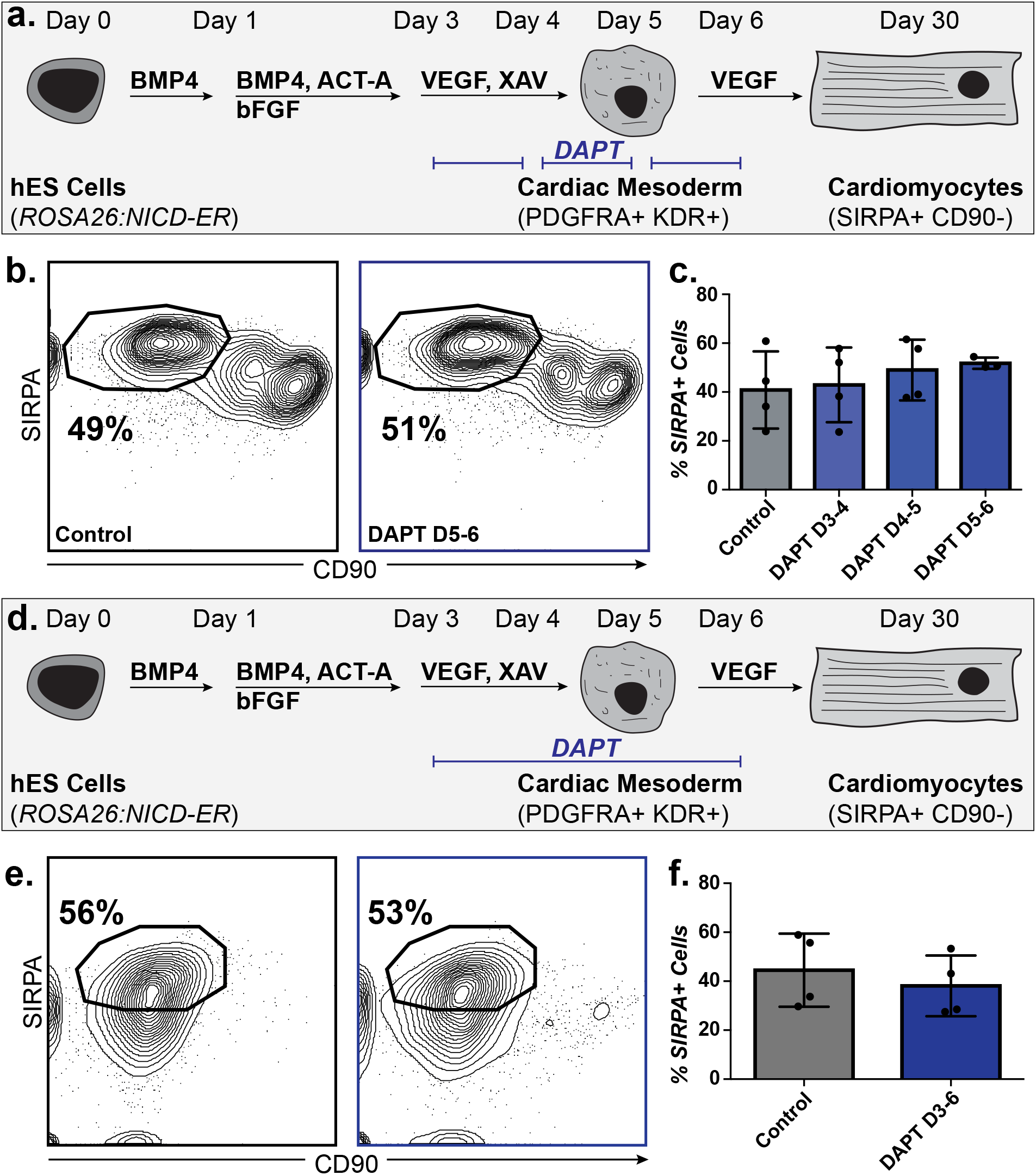
Inhibition of Notch has no effect on directed differentiation of cardiomyocytes from hPSCs. (a) Schematic of *ROSA26:NICD-ER* hPSC-cardiomyocyte differentiation and Notch inhibition times. (b) Flow cytometric analysis with antibodies against SIRPA and CD90 to measure cardiomyocyte (SIRPA+CD90-) differentiation efficiency at day 30 in control cultures (left) or after DAPT treatment from day 5-6 (right). (c) Quantification of data in (b). (d) Schematic of *MYL2-GFP* hiPSC-cardiomyocyte differentiation and Notch inhibition times. (e) Flow cytometric analysis with antibodies against SIRPA and CD90 to measure cardiomyocyte (SIRPA+CD90-) differentiation efficiency at day 30 in control cultures (left) or after DAPT treatment from day 3-6 (right). (f) Quantification of data in (e).

**Supplementary Figure 5:**
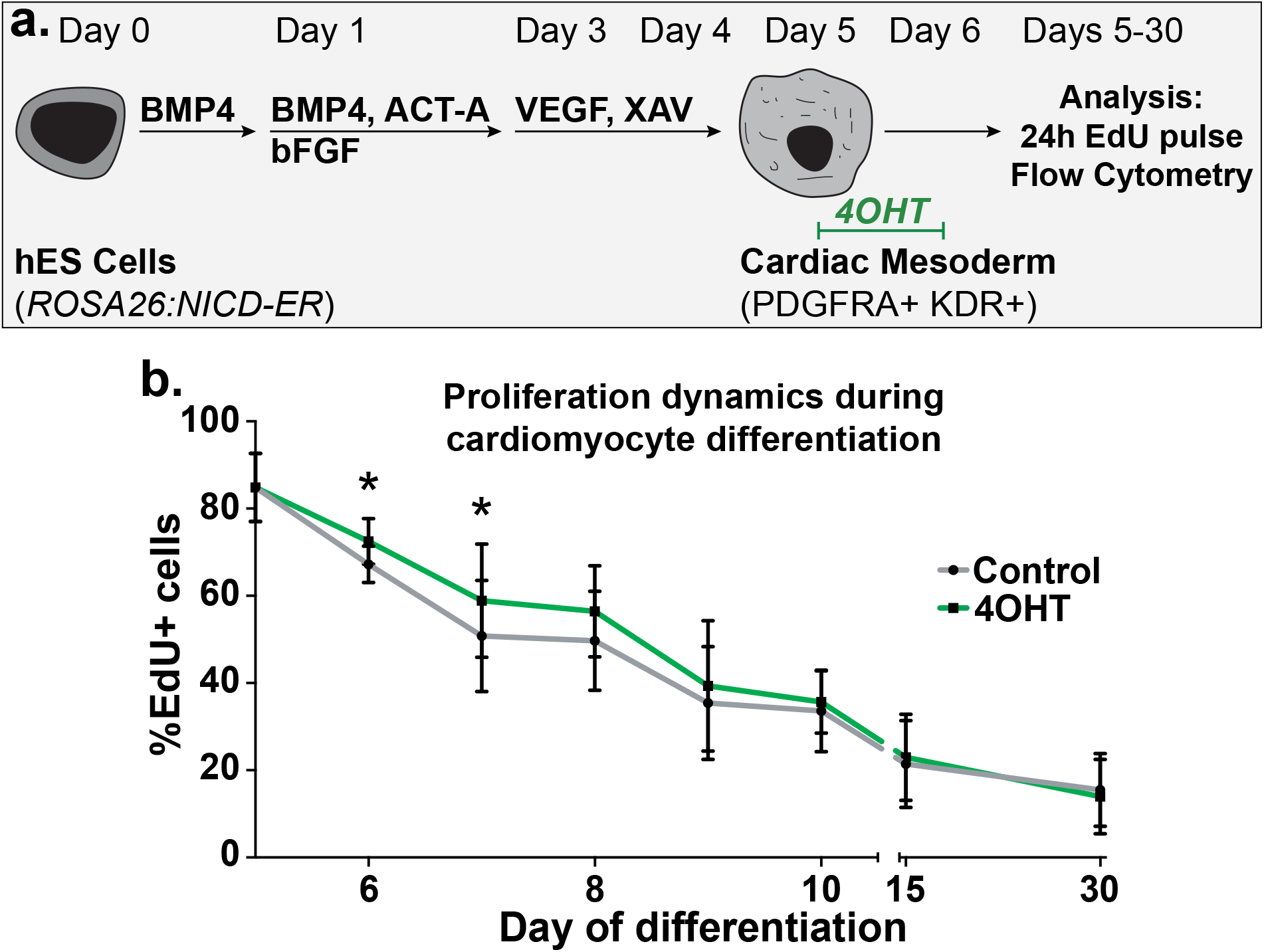
Notch induction briefly extends the proliferative window of cardiac precursors during hPSC directed differentiation. (a) Schematic of *ROSA26:NICD-ER* hPSC-cardiomyocyte differentiation and EdU analysis. Cultures were either untreated or treated with 4OHT from day 5-6 to induce Notch signaling. EdU was added 24h before each collection and analysis time. (b) EdU incorporation analysis by flow cytometry from day 5 to day 30 in control or Notch-induced cultures (n=5 differentiations).

**Supplementary Figure 6:**
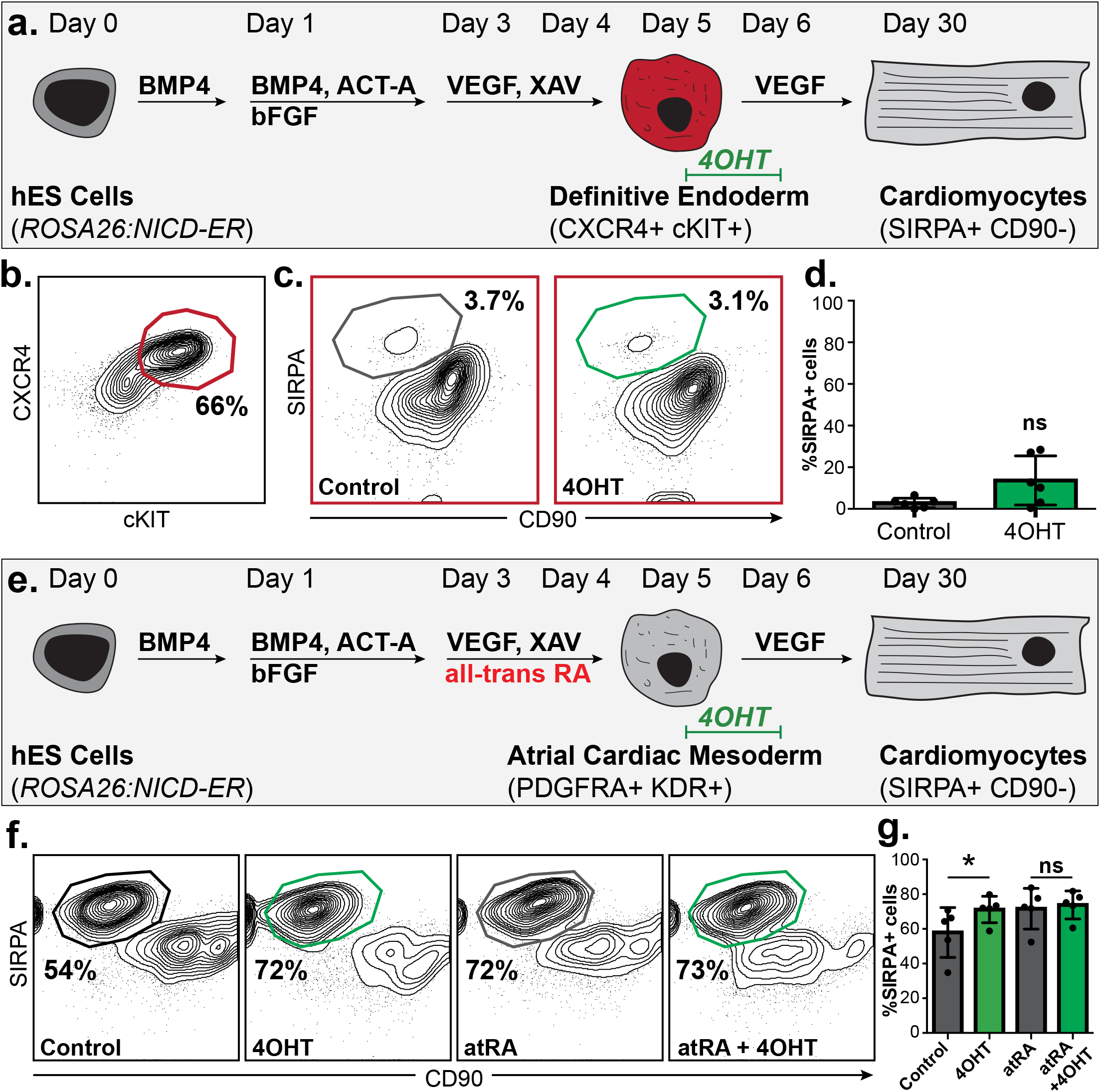
Induction of Notch signaling does not induce cardiomyocyte differentiation from definitive endoderm progenitors. (a) Schematic of *ROSA26:NICD-ER* hPSC differentiation to definitive endoderm progenitors, and continued differentiation in basal media. (b) Flow cytometric analysis of definitive endoderm progenitor generation with antibodies against CXCR4 and cKIT. (c) Flow cytometric analysis of cardiomyocyte generation from definitive endoderm cultured in basal media and the absence or presence of Notch induction from day 5-6. (d) Quantification of data in (c). (e) Schematic of *ROSA26:NICD-ER* hPSC differentiation to atrial cardiomyocytes through addition of all-trans retinoic acid (atRA). (f) Flow cytometric analysis of cardiomyocyte differentiation efficiency from ventricular (control) or atrial (atRA) cultures in the absence or presence of Notch induction from day 5-6. (g) Quantification of data in (f).

**Supplementary Figure 7:**
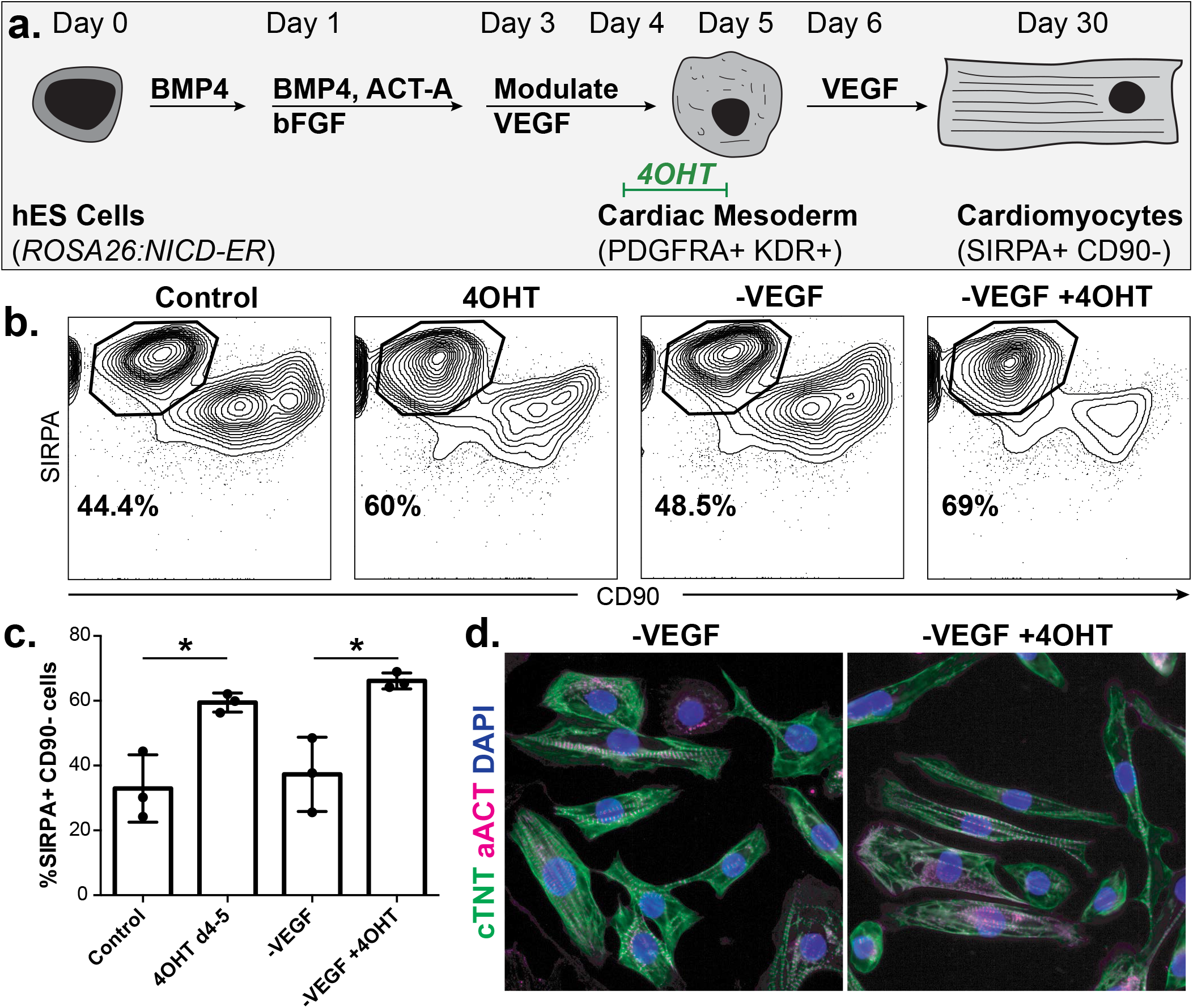
VEGF is not required for cardiomyocyte differentiation efficiency. (a) Schematic of *ROSA26:NICD-ER* hPSC-cardiomyocyte differentiation with WNT and VEGF modulation and Notch induction. (b) Flow cytometric analysis of cardiomyocyte generation in the absence of VEGF or WNT and VEGF, with and without Notch induction. (c) Quantification of data in (b). (d) Immunofluorescence analysis on day 30 cardiomyocytes with antibodies against cTNT and aACT.

